# Altered sensitivity to motion of area MT neurons following long-term V1 lesions

**DOI:** 10.1101/226571

**Authors:** Maureen A. Hagan, Tristan A. Chaplin, Krystel R. Huxlin, Marcello G. P. Rosa, Leo L. Lui

## Abstract

The middle temporal area (MT) receives its main afferents from the striate cortex (V1). However, MT also receives direct thalamic projections, which have been hypothesized to play a crucial role in residual vision after V1 lesions. MT neurons continue to respond shortly after V1 lesions, but human clinical work has shown that lesion effects can take up to six months to stabilize, making it important to understand MT responses after long-term deprivation of V1 inputs. We recorded neuronal responses in MT to moving dot stimuli in adult marmoset monkeys, 7-11 months after unilateral V1 lesions. Fewer MT neurons were direction selective, including neurons whose locations corresponded to the intact parts of V1. Firing rates were higher and more variable, and increased with motion strength regardless of direction. These properties could be re-created by a network model with imbalanced inhibition and excitation, providing the first insights into functional implications of long-term plasticity in MT following V1 lesions.

## Introduction

Damage to the striate cortex (V1; primary visual cortex) in adult primates leads to loss of conscious visual perception, referred to as cortical blindness (Lister and Holmes, 1916). In cases of partial destruction, a defined scotoma (an “island” of blindness) is created, which precisely follows the topographic representation of the visual field in V1 (Horton and Hoyt, 1991). Despite the absence of visual sensation, humans and monkeys with V1 lesions sustained in adulthood often retain a residual ability, termed “blindsight”, to respond to moving or flickering visual stimuli within the scotomas (Riddoch, 1917; Klüver, 1936; 1941; Poppel et al., 1973; Sanders et al., 1974; Weiskrantz et al., 1974; Barbur et al., 1993; Weiskrantz, 1996).

The middle temporal area (MT) of extrastriate cortex is a likely neural substrate for mediating the residual visual motion processing in blindsight. Within weeks of V1 lesions, many neurons in MT still respond in a direction selective way to oriented bars and gratings presented inside the scotoma (Rodman et al., 1989; Rosa et al., 2000; Azzopardi et al., 2003; Alexander and Cowey, 2008). However, humans with V1 damage exhibit the greatest rates of visual recovery, without therapeutic intervention, within the first 8 weeks after their lesion; once patients reach 6 months post-V1 damage, they no longer exhibit these improvements (Zhang et al., 2006). This suggests a change in neuronal responsiveness within spared visual cortices up to 6 months post-damage. Therefore, characterizing neuronal responses beyond 6 months is critical for understanding how plasticity in the adult visual system may allow for residual vision.

In patients with long-term V1 damage, the ability to determine the direction of complex motion stimuli, such as moving random dot patterns is extremely poor (Azzopardi and Cowey, 2001; Huxlin et al., 2009; Das et al., 2014; Cavanaugh et al., 2015). MT neurons in normal animals respond to moving random dot patterns in a way that directly reflects the animal’s psychophysical performance (Britten et al., 1992). The behavioral data from patients suggest that responses of MT neurons inside the lesion projection zone (LPZ; the region of cortex that originally received afferent connections from V1-damaged tissue) must be degraded after V1 lesions. However, recent studies found that with perceptual training, patients with V1 lesions can recover the ability to discriminate the direction of moving dot patterns (Huxlin et al., 2009; Das et al., 2014). This suggests that some motion encoding abilities in MT may also persist beyond 6 months post lesion, which can be recruited by directed training. However, the ability of MT neurons to encode motion after long-term V1 lesions remains unknown.

Several lines of evidence suggest functional responses of MT neurons may change due to different types spontaneous plasticity in the months following V1 lesions. Magnetic resonance imaging and tractography studies in human patients have found that white matter tracts between the lateral geniculate nucleus and MT (Ajina et al., 2015) and corpus callosum (Celeghin et al., 2017) adapt to compensate after loss of V1 input. In animal models of brain damage, loss of key sensory inputs has been shown to decrease the number inhibitory interneuron synapses in the LPZ (Keck et al., 2011), and to strengthen excitatory lateral connections, both within and outside the LPZ (Darian-Smith and Gilbert, 1994; Palagina et al., 2009; Yamahachi et al., 2009; Barnes et al., 2017). However, whether such mechanisms affect the responses of MT neurons has not been investigated.

To address these questions, we recorded the response properties of MT neurons to random dot stimuli in marmoset monkeys 7-11 months following V1 lesions. We confirm the prediction that MT neurons still respond to random dot patterns long after V1 lesions, although the prevalence of direction selectivity was significantly reduced and the sensitivity to motion coherence was significantly altered. By developing a network model of MT, we were able to demonstrate that the observed changes in neural responses were consistent with underlying changes in inhibition and lateral connectivity.

## Results

### Visual responsiveness and directional selectivity in MT after V1 lesions

The caudal portion of V1 was resected in four adult marmosets (Fig 1A-C, see Methods), which resulted in a corresponding LPZ in MT (Fig 1A,D-G) and scotoma in the visual field (Fig 1 inset, D-G). The extent of each scotoma was first determined by mapping the receptive fields of neurons on the edge of the remaining parts of V1 in all four animals with V1 resections (Fig 1D-G). The scotoma was defined as the area of the visual field in which V1 receptive fields could no longer be recorded (Fig. 1D-G; dashed outlines). The scotomas were elongated in shape, encompassed the central visual field, and extended to approximately 40° eccentricity in the right visual hemifield (Fig 1D-G).

**Figure 1.**
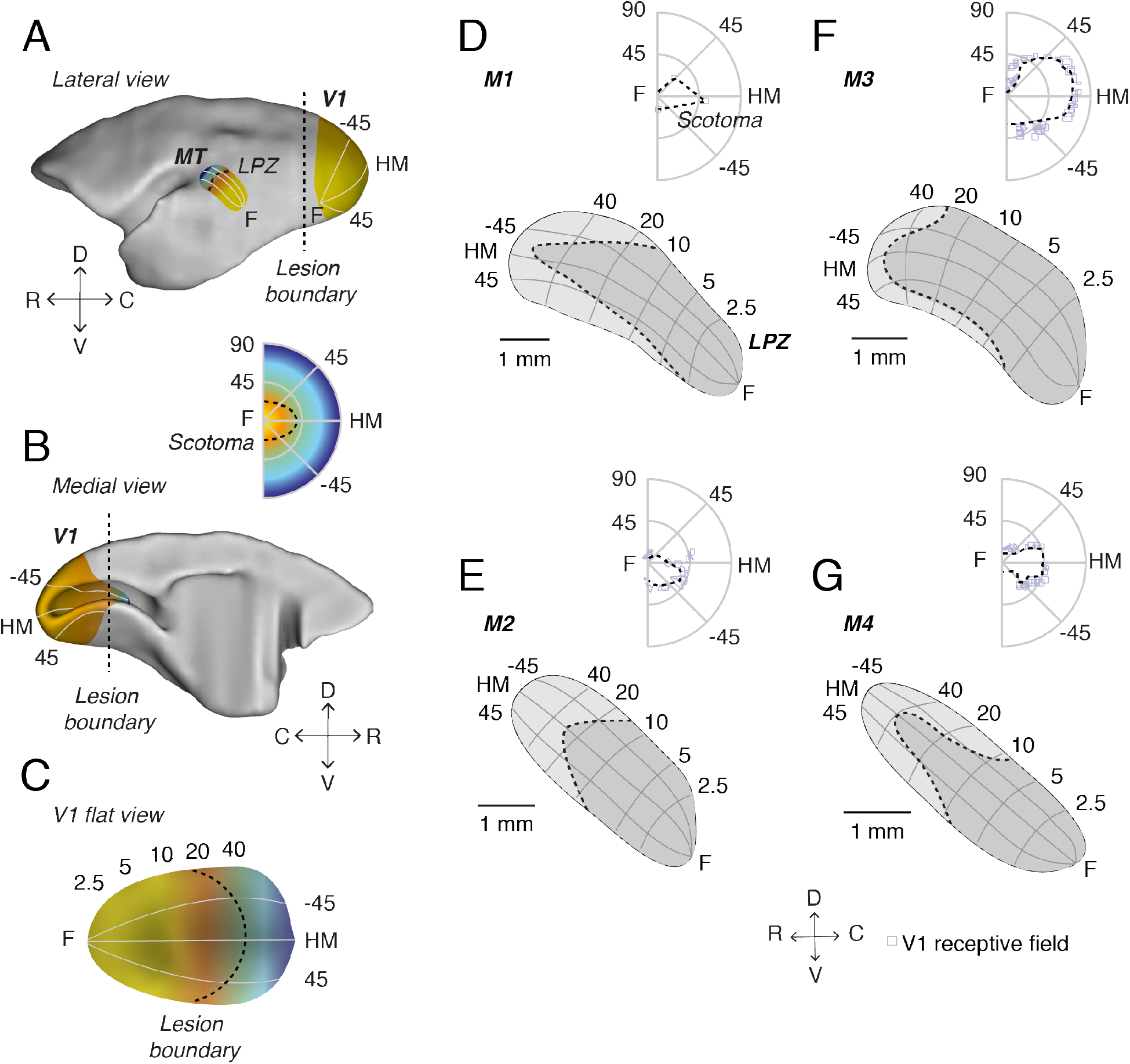
V1 lesions in adult marmosets. A, Lateral and B, medial views of a marmoset brain showing the locations of V1 and MT, as well as a C, flat view of V1. Illustrative retinotopic maps have foveal responses in yellow and peripheral responses in blue. Dashed lines illustrate the approximate location of the lesion boundary in V1 (A-C), and the corresponding LPZ in MT (A) and scotoma in the visual field (inset). D-G, V1 recordings were used to define the boundary of the scotoma in each V1-damaged animal and the scotoma was then projected onto the cortical surface representation of MT (see Methods) in order to determine the LPZ for each case. Numbers (degrees) indicate eccentricity (vertical axis) and polar angle (radial axes). The dashed lines indicate LPZ in MT and scotoma in visual field. C, caudal; D, dorsal; R, rostral; V, ventral; F, fovea; HM, horizontal meridian.

We recorded 114 single units and 134 multi-units from MT in four V1-damaged animals, and 36 single units and 38 multi-units (hereafter ‘units’) from two control animals. In all four V1-damaged animals, we found direction selective units in MT, both inside and outside the LPZ (Fig 2A-D). However, many units were not direction selective, and others showed no response to any of the stimuli we used. The majority of MT units in V1-damaged animals, both inside (73.3%, 115/157 units) and outside (79.1%, 72/91 units) the LPZ, responded to moving dot patterns. However, the percentage of visually-responsive units was significantly lower than in the control animals (Control: 97.3%, 72/74 units; compared to: Inside LPZ, p = 1.10x10^-21^; Outside LPZ, p = 9.91x10^-9^, Binomial distribution). Units with no visual response were excluded from further analysis. Consistent with previous studies (Eysel and Schweigart, 1999; Eysel et al., 1999; Rosa et al., 2000), we found that the majority of units (85.2%, 98/115 units) located inside the LPZ (which were expected to have receptive fields inside the scotomas according to the normal retinotopic organization of MT) had receptive fields displaced to the borders of the scotoma (Fig 2A-D).

**Figure 2.**
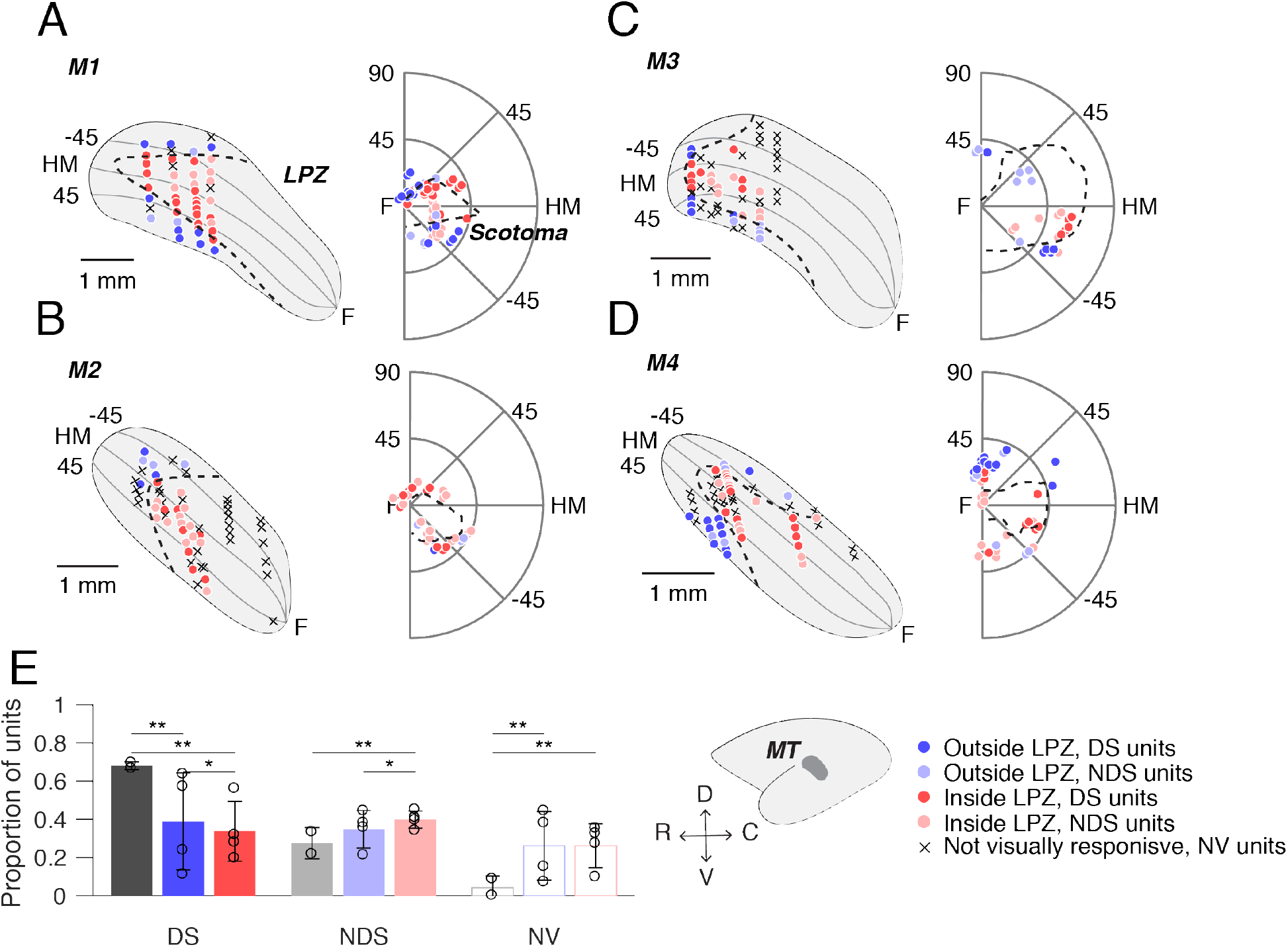
Direction selectivity in MT after V1 lesions. A-D, Anatomical locations of units recorded from MT and their respective receptive fields locations in the visual field in four adult marmosets (M1-M4, respectively) with V1 lesions. Each point indicates one unit in MT or one stimulus center. Dark colors indicate direction selective units (DS), light circles indicate units that were visually responsive but not direction selective (NDS). Red points indicate units recorded inside the LPZ and blue points indicate units outside the LPZ. Crosses indicate units that were not visually responsive (NV). Dashed lines indicate the LPZ in MT and the scotoma in the visual field. F, fovea; HM, horizontal meridian. Inset shows the position of MT on the cortical surface. E, Distribution of direction selective (DS), non-direction selective (NDS), and non-visually responsive (NV) units in control animals (grey) and in V1-damaged animals separated by whether the unit was located outside the LPZ (blue) or inside (red). Circles on each bar show percentages for individual subjects. Error bars indicate standard deviation. Stars indicate significance (one star: p < 0.05, two stars: p < 0.01, Binomial distribution).

We further divided responsive units based on direction selectivity. As expected, the majority of units in control animals were direction selective (Fig 2E, DS units: mean = 68.2%). However, significantly fewer MT units were direction selective in animals with a V1 lesion. This was true both outside (DS units: mean = 38.9%, p = 1.8x10^-5^, Binomial distribution) and inside (DS units: mean = 33.9%, p = 2.15x10^-19^) the LPZ, although the prevalence of direction selective units was higher outside than inside the LPZ (p = 0.02).

Although fewer MT units were direction selective in V1-damaged animals, there remained some highly direction selective units both outside (Fig 3A) and inside (Fig 3B) the LPZ, similar to controls (Fig 3C). To quantify the directional response in MT, we measured the direction selectivity index and circular variance. Units outside the LPZ in V1-damaged animals did not have significantly lower direction selectivity indices (Fig 3D, median = 0.74) in comparison with those from control animals (Fig 3F, Control, median = 0.81, p = 0.78, Wilcoxon’s rank sum). In comparison, units inside the LPZ had a significantly different distribution of direction indices, which was biased towards low values (Fig 3E, median = 0.65, p = 0.02). Likewise, the circular variance was significantly higher, indicating reduced direction selectivity, for units inside the LPZ compared to units outside the LPZ (Fig 3G, Outside LPZ: median = 0.66; Fig 3H, Inside LPZ: median = 0.75, p = 1.94x10^-3^) as well as to control units (Fig 3I, median = 0.61, p = 4.83x10^-5^). The circular variance of units outside the LPZ was not significantly different than controls (p = 0.27). Therefore, direction selectivity was reduced for units located inside the LPZ.

**Figure 3.**
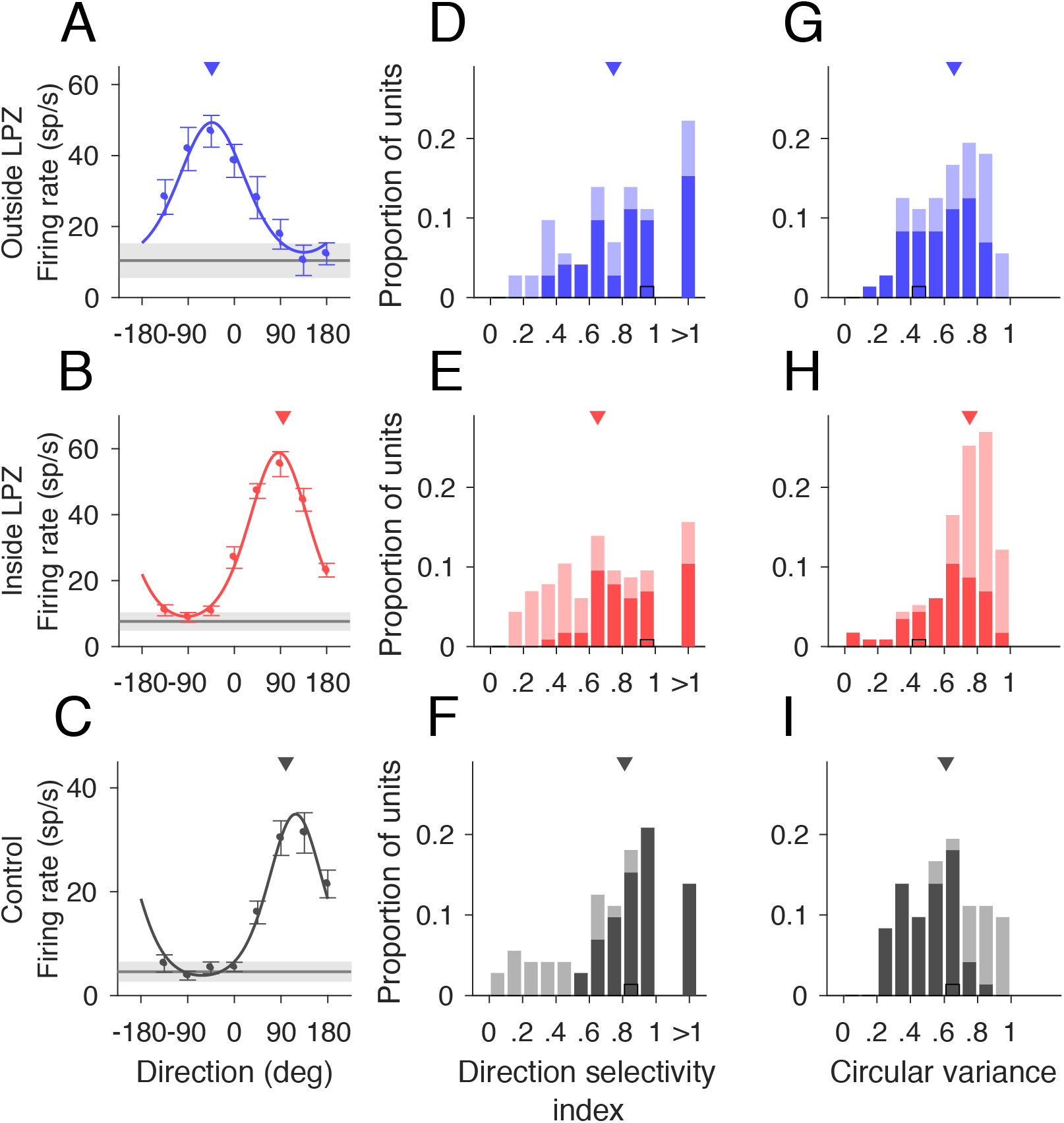
MT units inside LPZ are less direction selective. A-C, Three examples of direction selective units outside the LPZ, inside the LPZ, and in a control animal, respectively. Error bars indicate standard error, lines indicate fit of von Mises distribution, triangles indicate preferred direction. Grey line indicates mean spontaneous firing rate, shaded region indicates standard error. D-F, Direction selectivity index and G-I, Circular variance for units located outside the LPZ (n = 72 units), inside the LPZ (n = 115 units), and for the two control animals (n = 72 units), respectively. Darker bars indicate direction selective units, lighter bars indicate non-direction selective units. Black outlines indicate example units shown in A-C. Triangles indicate median of distribution.

### Impact of motion coherence on MT responses after V1 lesions

For typical direction selective MT units, firing rates increase with motion strength (i.e. coherence) in the preferred direction and decrease with motion strength in the null direction (e.g., Fig 4C, (Britten et al., 1992; Chaplin et al., 2017)). After V1 lesions, the majority of MT units tested for sensitivity to motion coherence, both direction selective and non-direction selective, did not show the expected monotonic relationship between firing rate and coherence. Instead, units both outside (Fig 4A) and inside (Fig 4B) the LPZ showed a “U-shaped” response to changes in motion coherence, meaning that firing rates were highest for motion at 100% coherence, regardless of stimulus direction.

**Figure 4.**
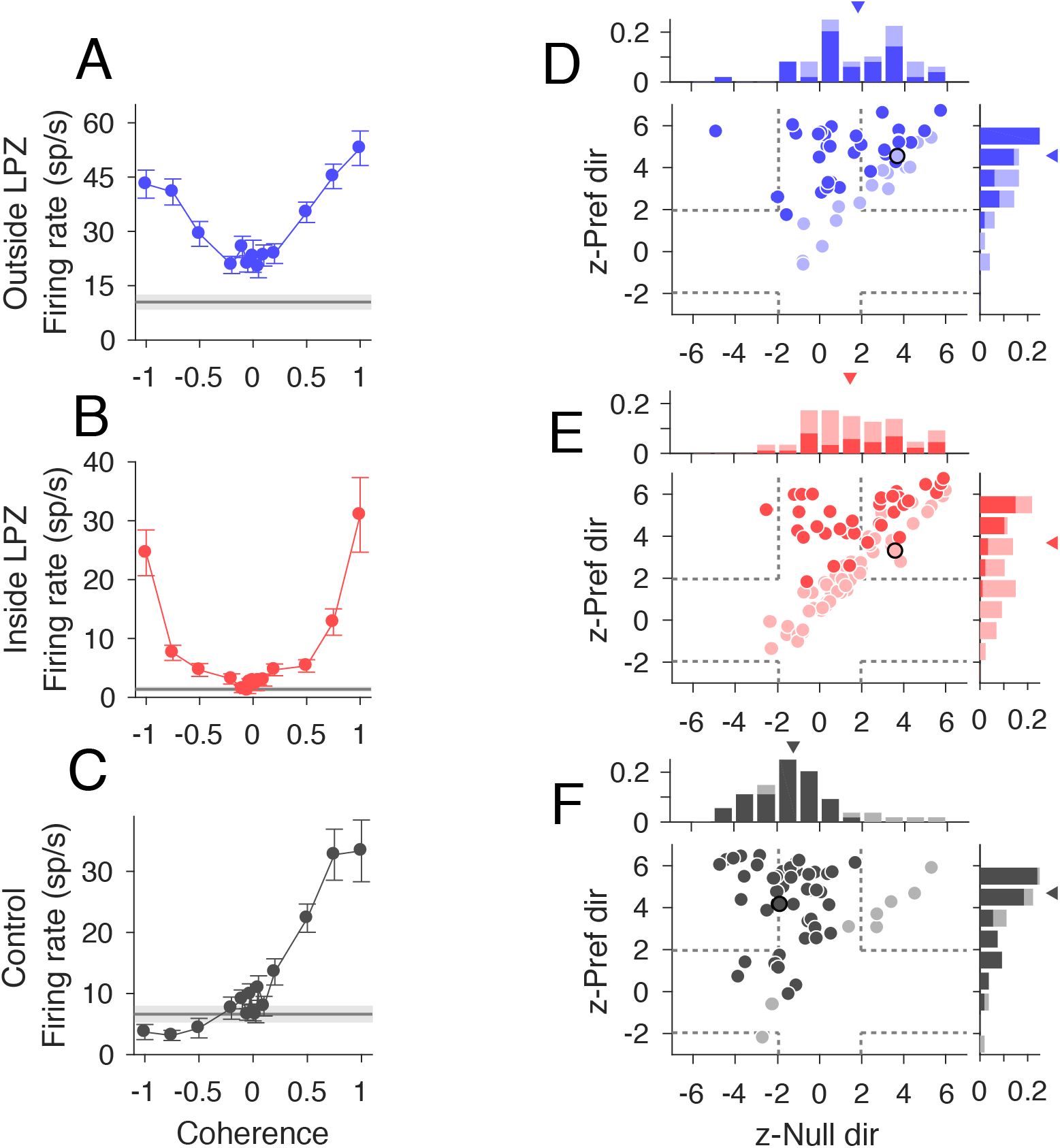
Responses to motion coherence after V1 lesions. A-C, Neural responses from three example units (outside the scotoma, inside the scotoma, and from a control animal, respectively) that showed typical responses for each population to motion coherence. Stimulus motion in the preferred direction is indicated by positive coherence values and in the null direction by negative coherence values. Average spontaneous firing rate indicated by the grey line; shaded area indicates standard error. D-F Scatter plots of z-scored firing rates for units located outside the LPZ (n = 49 units), inside the LPZ (n = 87 units), and for the control animals (n = 54 units), respectively. The y-axis shows the z-score for the firing rate difference between a stimulus with 100% coherence in the preferred direction and a stimulus with 0% coherence. The x-axis shows the z-score for the firing rate difference between a stimulus with 100% coherence in the null direction and a stimulus with 0% coherence. Each circle represents one unit. Darker circles indicate direction selective units, lighter circles indicate non-direction selective units. Black outlines indicate example units shown in A-C; Dashed grey lines indicate the significance threshold for each z-score (+/-1.96). Histograms show the distribution of z-scores for each axis. Triangle indicates median z-scores.

To quantify this curve shape, we calculated the z-score between firing rates at 100% coherence and 0% coherence for the preferred and null direction. Units with U-shaped responses had significantly positive z-scores (<1.96) for both the preferred and null direction (Top right quadrant of Fig 4D-F). Units with traditional linear responses had significantly positive z-scores in the preferred direction and negative z-scores in the null direction (Top left quadrant of Fig 4D-F). The majority of units both outside the LPZ (Fig 4D, 43/49, 87.5%, median = 4.53, p = 0.82, Wilcoxon’s rank sum, compared to controls) and inside the LPZ (Fig 4E, 57/87, 65.5%, median = 3.68, p = 0.02) showed significant increases in firing rate to the 100% coherent stimulus in the preferred direction, like control animals (Fig 4F, 44/54, 81.5%, median = 4.69). However, unlike controls, a significant percentage of units in V1-damaged animals also showed positive z-scores in the null direction (Outside LPZ: 24/49, 48.9%, median = 1.79, p = 8.47x10^-6^; Inside LPZ: 38/87, 43.6%, median = 1.43, p = 1.08x10^-6^; Control: 5/54, 9.3%, median = -1.25). Notably, this was true for both direction selective and non-direction selective units. Therefore, the majority of MT units in V1-damaged animals were sensitive to motion coherence, regardless of stimulus direction.

### Sensitivity to stimulus noise is impaired after V1 lesions

In macaques, it has been shown that the sensitivity of individual MT units is comparable to human behavioral performance when discriminating direction of noisy, random dot stimuli (Britten et al., 1992). Here, we asked whether the responses of MT units in chronic, V1-damaged animals are consistent with abnormal processing of stimulus noise, as was shown in humans with chronic V1 lesions (Azzopardi and Cowey, 2001; Huxlin et al., 2009; Das et al., 2014). We quantified the information carried by the firing rates of each unit by fitting a neurometric function and defining a direction threshold which was the coherence at which the performance was 0.82 (see Methods). Significantly fewer units in V1-damaged animals had direction thresholds (Fig 5A, example unit; Outside LPZ: 19/49, 38.8%, p = 1.22x10^-4^, Fig 5B, example unit; Inside LPZ: 23/87, 26.4%, p = 2.77x10^-13^, Binomial distribution) compared to control animals (Fig 5C, example unit; 35/54, 64.8%). Furthermore, the distribution of direction thresholds was significantly higher for units both outside the LPZ (Fig 5D; median = 0.80, p = 2.08x10^-3^, Wilcoxon’s rank sum) and inside the LPZ (Fig 5E, median = 0.79, p = 7.94x10^-3^) in comparison with controls (Fig 5F, median = 0.63). In summary, in line with the reduced direction selectivity, the sensitivity to direction in noise of MT units was substantially reduced in V1-damaged animals.

**Figure 5.**
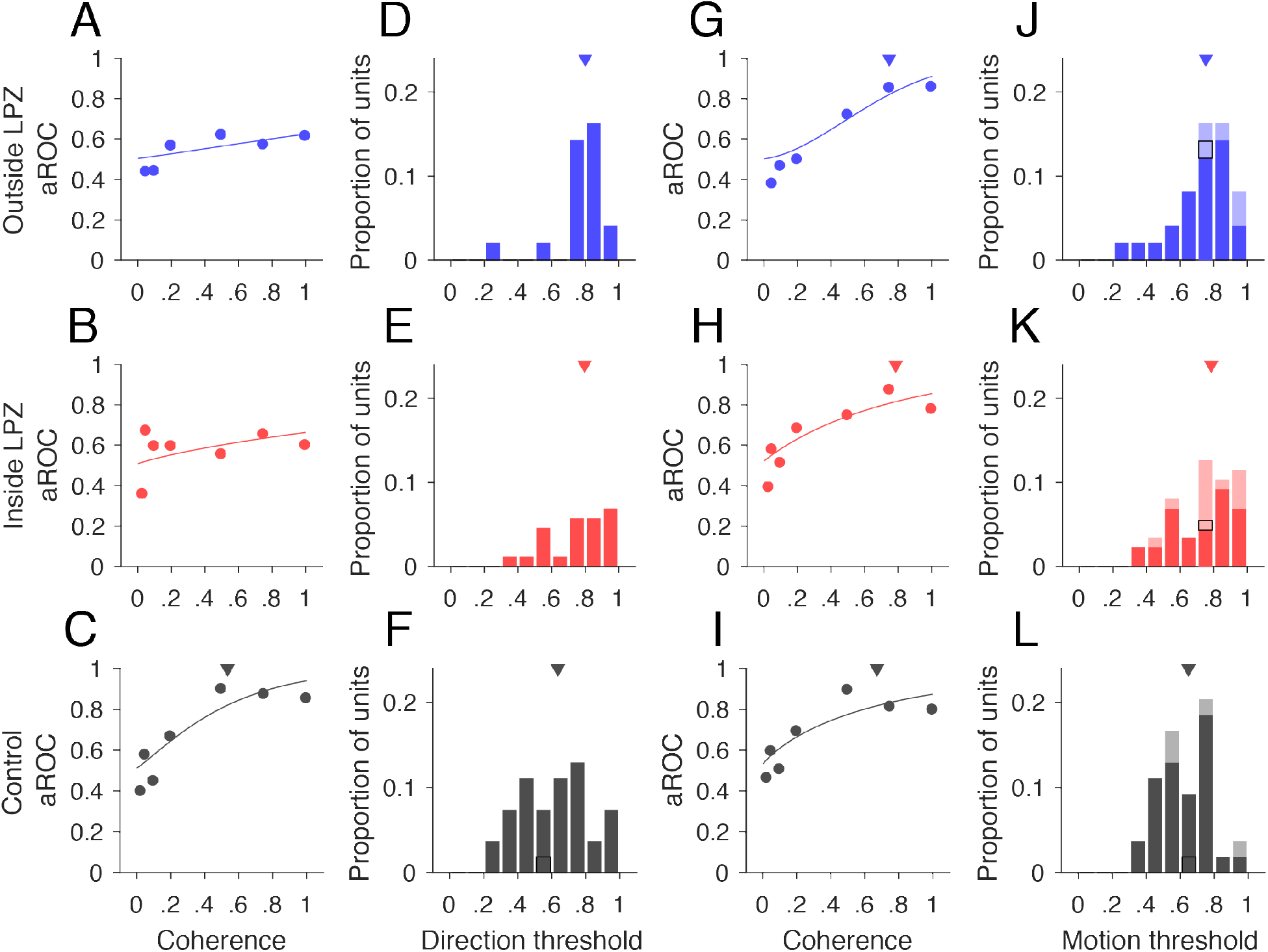
Direction thresholds and motion thresholds. A-C, Neurometric curves, data points indicate aROC values (preferred vs. null response) at each coherence for the same example units shown in Fig 5 A-C. The line indicates the best-fitting Weibull curve. Triangle in C shows the direction threshold coherence at which the curve reaches an aROC value of 0.82. D-F, Distributions of units with direction thresholds located outside the LPZ (n = 19/49 units), inside the LPZ (n = 23/87 units), and for the two control animals (n = 35/54 units), respectively. Black outlines indicate example units shown in A-C. Triangles indicate median threshold for each population. G-I, Neurometric curves, data points indicate aROC values (preferred vs. zero coherence response) at each coherence for same example units shown in Fig 5 A-C. All conventions as in A-C. J-L, Distributions of units with motion thresholds located outside the LPZ (n = 29/49 units), inside the LPZ (n = 45/87 units), and for the two control animals (n = 36/54 units), respectively. All conventions as in D-F.

Given that the majority of units in V1-damaged animals still showed significant increases in firing rate in response to increases in motion coherence, regardless of direction selectivity, we next compared the firing rate at each coherence in the preferred direction to the firing rate to the 0% coherence stimulus (Chaplin et al., 2017). This metric calculates the ability of the unit to detect motion, regardless of the direction. As for the direction thresholds, we defined the coherence at which the aROC curve reached 0.82 to be the unit’s motion threshold (Fig 5G-I). A greater percentage of units reached the motion threshold compared to the direction threshold, although for units with inside LPZ there were still significantly fewer that reached threshold compared to controls (Outside LPZ: 29/49, 59.2%, p = 0.06, Binomial distribution; Inside LPZ: 45/87, 51.7%, p = 1.37x10^-3^; Control: 36/54, 66.7%). The distributions of motion thresholds were also significantly higher than in controls for both units inside and outside the LPZ (Fig 6J, Outside LPZ: median = 0.75, p = 4.27x10^-3^, Wilcoxon’s rank sum; Fig 5K, Inside LPZ: median = 0.78, p = 3.91x10^-3^; Fig 5L, Control: median = 0.65). This indicates that the majority of units in animals with V1 lesions retain sensitivity to the strength of global motion, regardless of whether the unit is sensitive to the direction of motion.

**Figure 6.**
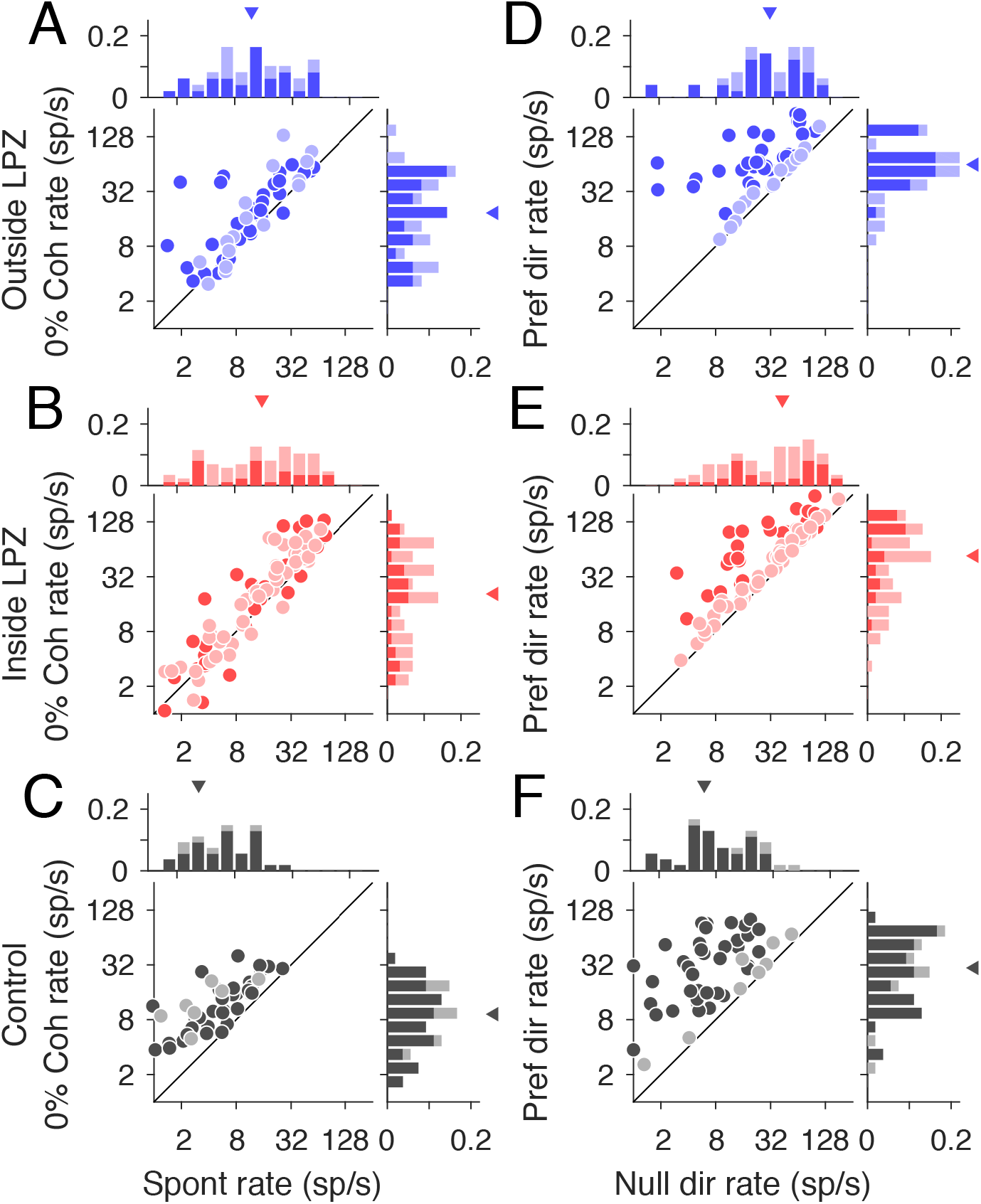
Firing rate differences in V1-damaged and control animals. A-C Spontaneous firing rates (x-axis) compared to firing rates for the 0% coherence stimulus (y-axis) for units located outside the LPZ (n = 49 units), inside the LPZ (n = 87 units), and for the two control animals (n = 54 units), respectively. D-F, Firing rates in the null direction (x-axis) compared to preferred (y-axis). Darker circles indicate direction selective units, lighter circles indicate non-direction selective units. Histograms show the distribution of firing rates for each axis. Darker bars indicate direction selective units, lighter bars indicate non-direction selective units. Triangles indicate median rate.

### MT firing rates are higher and more variable after V1 lesions

Metrics of direction selectivity and motion selectivity are dependent on the firing rates of units (Chaplin et al., 2017). If loss of V1 input reduces the overall firing rate of units inside the LPZ, this factor could account for the reduced prevalence of selectivity in MT. In fact, we found that firing rates were significantly higher in V1-damaged animals, compared to controls. Spontaneous rates were significantly higher outside the LPZ (Fig 6A, median spontaneous rate = 11.9 sp/s, p = 6.72x10^-8^) as well as inside the LPZ (Fig 6B, median spontaneous rate = 14.2 sp/s, p = 3.51x10^-9^; Fig 6C, Control: median spontaneous rate = 4.0 sp/s). Surprisingly, units in V1-damaged animals, as a population, did not have a significant visual response to stimulus with 0% motion coherence. In control animals, the response to a 0% coherence stimulus was typically above the spontaneous rate (median 0% rate = 8.8 sp/s; p = 4.32x10^-5^). In V1-damaged animals, this difference was not significant (Outside LPZ: median 0% rate = 20.1 sp/s, p = 0.07; Inside: median 0% rate = 21.2 sp/s, p = 0.17). This indicates that in V1-damaged animals, units were generally not visually responsive to stimuli that contained no net motion signal.

In addition to having higher spontaneous firing rates, MT units in V1-damaged animals had higher firing rates in the preferred and null directions as well (Fig 6D, Outside LPZ: median preferred rate = 63.5 sp/s, p = 3.28x10^-6^, median null rate = 32.5 sp/s, p = 7.07x10^-10^; Fig 6E, Inside LPZ: median preferred rate = 81.1 sp/s, p = 1.65x10^-4^, median null rate = 42.6 sp/s, p = 3.52x10^-11^, Wilcoxon’s rank sum), compared to units in control animals (Fig 6F, median preferred rate = 35.8 sp/s, median null rate = 5.8 sp/s). Therefore, for most units in V1-damaged animals, we did not find that the lack of direction selectivity or motion sensitivity was due to a decrease in the firing rate of units in V1-damaged animals. Rather, the reduced direction selectivity was driven by increased firing rates in the null direction.

In addition to analyzing the mean firing rate, we calculated the mean-matched Fano factor for single units across the trial (see Methods) to estimate whether variability in firing rate changes after loss of V1 input. While units still showed the characteristic dip in Fano factor at stimulus onset (Churchland et al., 2010), Fano factors remained high in the single units from V1-damaged animals. In fact, Fano factors were significantly higher in V1-damaged animals regardless of whether units were inside or outside the LPZ (Outside LPZ: Spontaneous median = 3.13, p = 1.0x10^-14^; Stimulus? median = 2.08, p = 6.65x10^-17^, Wilcoxon’s rank sum; Inside LPZ: Spont. median = 2.78, p = 2.03x10^-16^; Stim. median = 2.09, p = 1.91x10^-19^) compared to controls (Spont. median = 1.61; Stim. median = 1.25). In summary, firing rates were both higher and more variable in V1-damaged animals, which may reflect changes in the balance of local excitation (Litwin-Kumar and Doiron, 2012), as has been previously observed following loss of sensory input (Keck et al., 2011; Barnes et al., 2017).

### Modeling E-I imbalances inside the LPZ

Finally, to determine the extent to which altered MT responses may be driven by input into MT compared to local changes in the balance of excitation and inhibition (E-I), we developed a biophysical network model of MT. The model was composed of excitatory and inhibitory, leaky-integrate-and-fire neurons, containing two excitatory populations (Fig 8A, E1 and E2) that were tuned to opposite directions of motion. Excitatory and inhibitory populations were reciprocally connected (probability of connection = 0.2). Under balanced E-I conditions, E1 and E2 showed monotonic increases to changes in stimulus coherence in their respective preferred directions (Fig 8B, mean response for each population) and the spontaneous rates were low (mean rate = 1.91 sp/s), similar to MT responses in control animals. By modeling the stimulus as a time-varying direct input to each neuron, consisting of a visual component and direction selective component (see Methods), we were able to manipulate this direct input to be direction selective visual input, non-direction selective visual input, or non-visual input (Fig 8A). We were then able to create sub-populations of E1 and E2 that were “inside” or “outside” the LPZ by manipulating the input into those neurons. Half of the neurons in the pool were “outside” the LPZ; these always received the standard visual and direction selective stimulus inputs. The other half consisted of neurons “inside” the LPZ. We tested whether cells inside the LPZ that received direction selective visual input, non-direction selective visual input, or non-visual inputs best reproduced our observed data. We also compared balanced E-I connections to imbalanced E-I connections. In the latter condition, inhibitory synapses were reduced and excitatory lateral connections were strengthened, as has been observed experimentally (Darian-Smith and Gilbert, 1994; Palagina et al., 2009; Yamahachi et al., 2009; Keck et al., 2011). Because the responses of E1 and E2 were mirrored (Fig 8B), for clarity, only the results of population E2 (neurons that preferred the positive direction on the x-axes) are shown.

**Figure 7.**
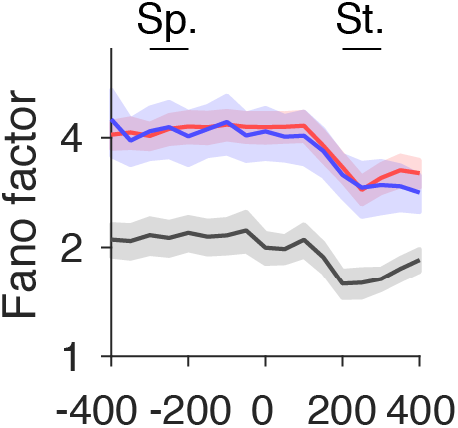
Mean-matched Fano factors in V1-damaged and control animals. Mean-matched Fano factors across the trial for units with receptive fields outside scotoma (blue, n = 124 conditions), inside scotoma (red, n = 256 conditions) and from control animals (grey, n = 192 conditions). Error bars indicate 95% confidence intervals. Sp.,spontaneous interval; St., stimulus interval.

**Figure 8.**
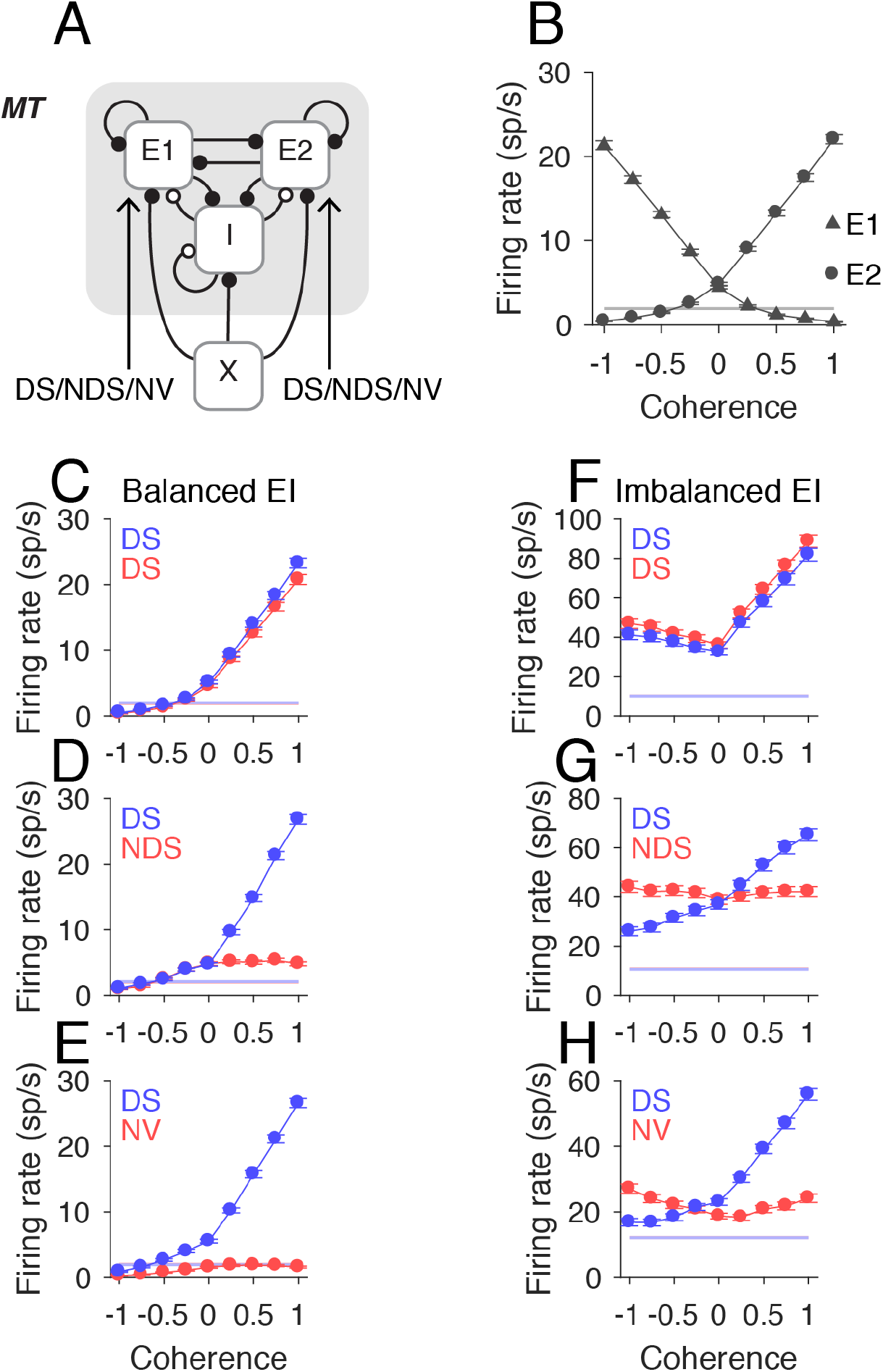
Network model of LPZ. A, Schematic of network model for an MT-like circuit composed of two populations of excitatory neurons (E1, E2) tuned to opposite directions of motion and reciprocally connected inhibitory (I) neurons. Each population receives an external Poisson signal (X). The excitatory populations also receive sensory input, which may be direction selective visual input (DS), non-direction selective visual input (NDS), or non-visual input (NV). B, Population mean firing rates for E1 neurons (triangles, n = 800) and E2 neurons (circles, n = 800) in response to varying motion coherence in two directions. Grey line indicates mean spontaneous rate, error bars indicate standard error. C-E, Population mean firing rates for E2 neurons “outside” the LPZ (blue circles, n = 400) and “inside” the LPZ (red circles, n = 400) in the balanced E-I network where the neurons “outside” the LPZ received direction selective input (DS) and the neurons inside the LPZ received either direction selective visual input (DS), non-direction selective visual input (NDS) or non-visual input (NV), respectively. Circles indicate mean population rate, error bars show standard error. F-G Population mean firing rates for E2 neurons in the imbalanced E-I network. All other conventions the same as C-E.

In the balanced E-I network, the spontaneous rates were unchanged from the control conditions (mean spontaneous rate = 2.01 sp/s; p = 0.27, Wilcoxon’s rank sum). The population response of neurons outside the LPZ maintained a consistent monotonic response to coherence, regardless of input into the LPZ (Fig 8C-E). Neurons inside the LPZ also showed a monotonic response to coherence when receiving direction selective inputs, as expected (Fig 8C), but showed a flat response when only receiving non-direction selective inputs (Fig 8D) or non-visual input (Fig 8E). This did not match the observations in the V1-damaged animals.

While maintaining the same inputs, we then manipulated the probability of connections in the network (see Methods) to create an E-I imbalance. As a result, the firing rates increased, including the spontaneous rates (mean spontaneous rate = 12.21 sp/s, = 0, Fig 8F-H), as observed experimentally (Fig 6).

Furthermore, when input into the LPZ was either direction selective (Fig 8F) or not visually responsive (Fig 8H), the responses to motion coherence in the null direction increased, as was observed experimentally, although this was only significant for non-visual inputs (Fig 8F, Inside DS, Null rate at 100% coherence vs 0% coherence, p = 0.48; Fig 8H, Inside NV, p = 0.02, Wilcoxon’s rank sum). In the case of direction selective input, this was true for neurons outside the LPZ as well, although this was also not significant (Fig 8F, Outside DS, p = 0.91). When the input into the LPZ was non-direction selective, the responses to motion coherence became flat (Fig 8G), as in the balanced network condition. Therefore, our modeling results suggest that the observed responses in MT may be due to an imbalance in excitation and inhibition caused by the V1 lesion, and that the input into the LPZ is likely to be a mix of direction selective and non-selective inputs.

## Discussion

In order to understand the long-term consequences of V1 lesions, we recorded responses in MT to moving random dot patterns in adult marmoset monkeys. More than 7 months after the lesions, we found that fewer units in MT were direction selective, regardless of whether they were located inside or outside the LPZ. At the same time, sensitivity to motion coherence was largely preserved, even for a large number of non-direction-selective units. Therefore, a considerable amount of motion processing is still carried out in MT, despite the decreased direction selectivity. Interestingly, the firing rates of the cells were higher and more variable in V1-damaged animals. These observations were predicted by a network model, in which the balance of excitatory and inhibitory connections within MT changed after chronic V1 lesions. Together, these results provide novel insights to the underlying mechanisms of neural changes after chronic V1 damage.

### Motion selectivity in MT after V1 lesions

Consistent with previous work (Rodman et al., 1989; Rosa et al., 2000; Azzopardi et al., 2003), our data confirmed the preservation of a group of direction-selective MT units inside the LPZ. However, we found fewer direction-selective cells in MT compared to previous studies, although the latter were done at shorter time scales after V1 lesions and used oriented bars and gratings as opposed to random dot patterns. Both time since damage and choice of stimulus may account for differences found in the present study. In addition, we observed robust (albeit abnormal) sensitivity to motion coherence, despite the reduced direction selectivity. The increased direction thresholds for noisy moving dot stimuli were also indicative of reduced direction selectivity. These findings are consistent with the notion that direction selectivity in MT may be generated in V1 (Movshon and Newsome, 1996).

Surprisingly, we found decreased direction selectivity and abnormal sensitivity to motion coherence in neurons located both inside and outside the LPZ. Previous work in adult animals with short-term V1 damage had found neural responses in the region outside the LPZ to be largely preserved, using bar and square-wave grating stimuli (Rosa et al., 2000). Because MT receptive fields have a finite size (Rosa and Elston, 1998), it is likely that neurons near the estimated boundary of the LPZ receive different degrees of input from the remaining parts of V1. Likewise, the receptive fields of cells inside the LPZ tended to be closer to and often overlapped the scotoma border, as has been previously reported following cortical (Rosa et al., 2000; Giannikopoulos and Eysel, 2006) and retinal lesions (Schmid et al., 1996). Thus, the surviving responses inside the LPZ in MT may be dependent on proximity to units outside the LPZ, which still receive inputs from V1. Together, these results are indicative of potentially widespread changes in the connectivity of units in MT, or at least a substantial change in the balance of inputs previously present, both inside and outside the LPZ, after V1 lesions.

### MT connectivity after V1 lesions

In addition to inputs from V1 and extrastriate cortical areas, MT also receives direct subcortical input from the koniocellular layers of the lateral geniculate nucleus in both macaques (Sincich et al., 2004) and marmosets (Warner et al., 2010). Many neurons in these layers have “cortical like” response properties (White et al., 2001; Solomon et al., 2010), including direction selectivity (Cheong et al., 2013), although these properties may be dependent on cortical feedback. MT also receives input from the inferior pulvinar nucleus (Berman and Wurtz, 2010; 2011). Both the lateral geniculate nucleus (Bridge et al., 2010; Schmid et al., 2011; Ajina et al., 2015) and the pulvinar (Rodman et al., 1990; Warner et al., 2012; 2015) have been implicated in the preservation of MT responses after V1 damage. Many neurons in the LPZ of the lateral geniculate nucleus degenerate over the first several months after V1 lesions (Atapour et al., 2017), whereas the projections to MT appear to be preserved (Bridge et al., 2008; Ajina et al., 2015). Together, these results suggest that subcortical inputs to MT may also contribute to residual responses in MT after V1 lesions, particularly for responses observed well inside the scotoma.

It is also possible that changes in local connectivity occur following V1 lesions. Shorter-term lesions and temporary inactivation studies of V1 have reported lower firing rates in MT, particularly for units inside the LPZ (Rodman et al., 1989; Girard et al., 1992). Contrary to this, we found higher than normal firing rates (both spontaneous and stimulus-induced) in MT more than 6 months post-V1 damage. We also found that the Fano factors for MT single units in V1-damaged animals were significantly higher than those of controls. Previous modeling work has suggested that high Fano factors can be achieved via “over clustering” of excitatory responses (Litwin-Kumar and Doiron, 2012). It may be the case that after V1 lesions, the units that remain active become more inter-connected with one another, leading to higher variability and broader directional tuning. Previous studies have found that a lack of sensory input decreases the number of inhibitory interneuron synapses (Keck et al., 2011), and hyper-excitability has been observed following retinal lesions (Giannikopoulos and Eysel, 2006). Furthermore, strengthening of excitatory lateral connections from within the LPZ and from neighboring cortical areas has been observed after loss of sensory input (Darian-Smith and Gilbert, 1994; Palagina et al., 2009; Yamahachi et al., 2009). Consistent with this, our network model suggests that an E-I imbalance, driven by increased excitability of lateral interactions and decreased probability of inhibitory synapses, could explain the observed decreased direction selectivity, “U-shaped” response to motion coherence and increased firing rates, independent of any additional input into the LPZ of MT. Neurons located near the LPZ boundary may still receive a greater proportion of direction-selective input from V1 than neurons further from the boundary, but our model suggests that these responses would also become “U-shaped” under imbalanced E-I conditions.

### Implications for adult humans with V1 damage

Patients who suffer damage to V1 usually experience a period of spontaneous behavioral recovery, which may last up to six months (Zhang et al., 2006). The majority of hemianopic studies characterize behavioral performance after this period, when the visual field defect has stabilized. The results of the present study, performed more than 6 months post-lesion, provide the most meaningful comparisons to date between non-human primate MT physiology and the behavioral performance of hemianopic patients. The one previous study performed on a similar timescale found no visual responses inside the scotoma of New World monkeys (Collins et al., 2003), a result that may be linked to the use of different anesthetics (Girard et al., 1992). We also found a high percentage of units that were unresponsive to our stimulus, which would likely have had receptive fields inside the scotoma before the lesion, based on their anatomical location within MT.

Hemianopic patients cannot easily discriminate the direction of motion of random dot stimuli (Azzopardi and Cowey, 2001), a result that correlates well with the poor direction selectivity we observed in similar conditions. However, recent studies have found that perceptual training protocols can lead to improvements in global motion discrimination in cortically blind patients (Sahraie et al., 2006; Raninen et al., 2007; Huxlin et al., 2009; Sahraie et al., 2010; Das et al., 2014). For this to occur, there must be neurons with sufficient motion encoding that can be recruited. The robust sensitivity to motion coherence we find in our data and the few remaining direction selective cells suggest that MT is a likely site to mediate these perceptual improvements. The responses we recorded along the border of the scotoma are also consistent with the finding that functional recovery with perceptual training in patients with chronic V1 lesions occurs primarily at the edges of the scotoma (Cavanaugh and Huxlin, 2017).

Taken together, our findings demonstrate that the rudiments of global motion processing persist in area MT of marmosets with long-standing (>6 months) V1 lesions. The functional changes we observed are consistent with long-term changes in the structural inputs into MT neurons. These changes may provide the infrastructure for continued motion perception. Furthermore, our results suggest that MT is a likely neural substrate for the improvements in motion perception observed in patients with V1 damage. Therefore, one would predict that the sensitivity to random dot patterns of MT neurons would improve during perceptual training.

## Materials and Methods

### Animals and surgical preparation

Single and multi-unit extracellular recordings were obtained from 6 adult marmoset monkeys. Experiments were conducted in accordance with the Australian Code of Practice for the Care and Use of Animals for Scientific Purposes and all procedures were approved by the Monash University Animal Ethics Experimentation Committee. Four of the marmosets received V1 lesions (M1 to M4), and at time of lesion, they were aged > 2 years. Neural data from MT were collected from two additional adult marmosets (M5, M6) of comparable age at the time of the recordings, who did not receive V1 lesions.

### Cortical lesions

Intramuscular (i.m.) injections of atropine (0.2 mg/kg) and diazepam (2 mg/kg) were administered as premedication, 30 minutes prior to the induction of anesthesia with alfaxalone (Alfaxan, 10 mg/kg, Jurox, Rutherford, Australia). Dexamethasone (0.3 mg/kg i.m., David Bull, Melbourne Australia) and penicillin (Norocilin, 50 mg/kg, i.m.) were also administered. Body temperature, heart rate, and blood oxygenation (P0_2_) were continually monitored, and supplemental doses of anesthetic were administered when necessary to maintain areflexia. Under sterile conditions, a craniotomy was made over the occipital pole of the left hemisphere. Using a fine-tipped cautery, an excision was then made of all cortical tissue caudal to a plane extending from the dorsal surface of the occipital lobe to the cerebellar tentorium, across the entire mediolateral extent (Rosa et al., 2000). The caudal 6 – 8mm of cortex, approximately two-thirds of V1, was removed entirely, including the occipital operculum, the exposed medial and ventral surfaces, and the caudal part of the calcarine sulcus (Fig. 1A-C). After application of hemostatic microspheres, the exposed cortex and cerebellum were protected with ophthalmic film, and the cavity was filled with Gelfoam. The skull flap was repositioned and secured with cyanacrylate (Vetabond, 3M), and the skin was sutured. Throughout the post-lesion period, monkeys were housed in large cages in groups with access to both indoor and outdoor environments. Following the V1 lesion surgery, animals recovered for 7-11 months before electrophysiological recordings.

### Electrophysiological recordings

The preparation for electrophysiological studies of marmosets has been described previously (Yu et al., 2010), and only the main points will be summarized here. Injections of atropine (0.2 mg/kg, i.m.) and diazepam (2 mg/kg, i.m.) were administered as premedication, 30 minutes prior to the induction of anesthesia with alfaxalone (Alfaxan, 10 mg/kg, i.m., Jurox, Rutherford, Australia), allowing a tracheotomy, vein cannulation and craniotomy to be performed. After all surgical procedures were completed, the animal received an intravenous infusion of pancuronium bromide (0.1 mg/kg/h; Organon, Sydney, Australia) combined with sufentanil (6 μg/kg/h; Janssen-Cilag, Sydney, Australia) and dexamethasone (0.4 mg /kg / h; David Bull, Melbourne, Australia), and was artificially ventilated with a gaseous mixture of nitrous oxide and oxygen (7:3). The electrocardiogram and level of cortical spontaneous activity were continuously monitored. Administration of atropine (1%) and phenylephrine hydrochloride (10%) eye drops (Sigma Pharmaceuticals, Melbourne, Australia) resulted in mydriasis and cycloplegia. Appropriate focus and protection of the corneas from desiccation were achieved by means of hard contact lenses selected by streak retinoscopy.

Neural recordings were made using single parylene-coated tungsten microelectrodes (0.7-1.2 MΩ) with exposed tips of 10 μm (Mircoprobe, MD). Electrophysiological data were recorded using a Cereplex system (Blackrock Microsystems, MD) with a sampling rate of 30 kHz. Each channel was high-pass filtered at 750 Hz. Spike waveforms were extracted off-line. Spike waveforms were then over-clustered using principal component analysis and manually merged on a moving 100 ms time window. Any remaining threshold crossings were classified as multi-unit activity.

### Stimuli

Computer generated stimuli were presented on a Display ++ monitor (M1-M4; 1920 x 1080 pixels; 710 x 395 mm; 120 Hz refresh rate, Cambridge Research Systems) or a VIEWPixx3D monitor (M5-M6; 1920 x 1080 pixels; 520 x 295 mm; 120 Hz refresh rate, VPixx Technologies) positioned 0.35 to 0.45 m from the animal on an angle to accommodate the size and eccentricity of the receptive field(s). All stimuli were generated with MATLAB using Psychtoolbox-3 (Brainard, 1997; Pelli, 1997).

Stimuli for quantitative analysis consisted of random dots presented in circular apertures. Dots were white and displayed on a black background, and were 0.2 deg in diameter. The density was such that there were on average 0.5 dots per deg^2^. Dot coherence was controlled by randomly choosing a subset of “noise” dots on each frame which were displaced randomly within the stimulus aperture. The remaining “signal” dots were moved in the same direction with a fixed displacement.

### Determination of Scotoma

The location of the scotoma was determined by mapping the receptive fields of neurons on the edge of the remaining parts of V1 (Fig 1D-G). This was done either on the Display++ monitor using static flashed squares (M1) or by hand mapping with luminance defined stimuli on a hemispheric screen (Yu and Rosa, 2010). As reported elsewhere (Rosa et al., 2000; Yu et al., 2013), the locations of receptive fields of neurons located on both the dorsal and ventral banks of the calcarine sulcus were similar to those found at corresponding locations in normal animals (Fritsches and Rosa, 1996; Chaplin et al., 2013). The scotoma was defined as the area of the visual field in which V1 receptive fields could no longer be recorded (Fig. 1D-G; dashed outlines).

### Determination of MT receptive fields

In MT, receptive fields were quantitatively mapped using a grid of stimuli presented across the screen. These stimuli were either small apertures of briefly presented moving dots (300 ms, diameter 5 degrees, animals M1-M4) or static flashed squares (length 4 degrees, animals M5 and M6). For quantitative tests, stimuli were presented inside a circular aperture restricted to the excitatory receptive field.

### Quantitative Tests

We conducted a series of tests to determine direction selectivity, speed tuning and sensitivity to motion coherence. All stimuli were presented for 500 ms with an inter-stimulus interval of 1000 ms. Direction and speed tuning tests used at least 12 repeats of each stimulus type, and motion coherence tests used 25 repeats.

We tested for direction selectivity by presenting a circular aperture of random dots that moved in 1 of 8 directions (0, 45, 90, 135, 180, 225, 270, 315) at 8 deg/s. Speed tuning was then tested using random dot stimuli with speeds (0, 2, 4, 8, 16, 32, 64, 128 deg/s) moving in the preferred and null direction. Stimuli with different motion coherences (0, 5, 10, 20, 50, 75, 100 %) were presented at a near-preferred speed, in both preferred and null directions.

### Data Analysis

*Visual response*: Neurons were considered to be visually responsive if the mean stimulus-evoked activity across all conditions was significantly greater than the spontaneous rate (Wilcoxon rank sum test, p < 0.05).

*Direction selectivity:* The preferred direction was calculated using a vector sum of normalized above-spontaneous spiking rates (Ringach et al., 2002). In each task, neurons were classified as direction selective if the response in the preferred direction was significantly greater than the null direction (Wilcoxon rank sum test, p < 0.05). To measure the degree of direction selectivity, we used three metrics.

First, we calculated a direction selectivity index (DSI):

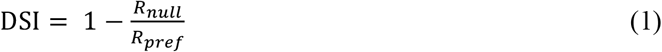

where R_pref_ and R_null_ are the above spontaneous spike rates in the preferred and null directions, respectively. Circular variance (CV) was calculated as 1 minus the length of the vector sum of normalized above-spontaneous spiking rates (Ringach et al., 2002). We also employed ideal observer analysis to determine the performance of MT neurons in discriminating two directions (Britten et al., 1992). This was calculated using the Receiver Operator Characteristic (aROC) curve for the distributions of responses to the preferred and null directions.

*z-Score firing rate differences*: We used random permutation to determine if the difference in mean between two conditions was significantly different than a distribution of mean differences from trials that were shuffled and chosen at random. We repeated this permutation 10,000 times to obtain a distribution.

*Direction thresholds:* In order to quantify a unit’s susceptibility to motion noise, we employed ideal observer analysis to determine performance of MT neurons in a direction discrimination task (Britten et al., 1992). For each level of coherence, aROC was calculated and the aROC values were fitted using least squares regression with a Weibull function, resulting in a neurometric curve that described the neuron’s performance with respect to coherence.

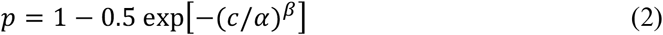

where *p* was the probability of correctly discriminating the direction of motion at coherence *c*, α was the coherence of threshold performance (p=0.82) and β controls slope. The α was limited between 0 and 3, and the β was limited to be between 0 and 10. Neurons that do not have an aROC of at least 0.82 at 100% coherence cannot have a threshold (i.e. p(c=100)<0.82), and were excluded from analyses of thresholds, as was any neuron whose threshold exceeded 100%.

*Motion Thresholds:* In order to determine how well neurons could detect motion in the preferred direction versus random motion, we calculated aROC comparing the distribution of spikes evoked by each level of motion coherence with the distribution of spikes evoked by zero coherence. As for direction thresholds, we fit a Weibull function (equation (2) above) to these data to determine the detection threshold.

*Mean-matched Fano factor:* To compute the variance in firing rates over time, we computed the mean-matched Fano factor using the procedures described by Churchland and colleagues (Churchland et al., 2010) using the Variance toolbox (Stanford University) in MATLAB. Briefly, for each single unit we used the firing rates from the motion coherence task. Each stimulus condition (motion coherence and direction) was treated separately. Spike counts were computed using a 100 ms sliding window moving in 50 ms time steps. To control for differences in firing rates, we computed a mean-matched Fano factor. For each time window, we matched the mean firing rates by randomly deleting spikes until mean rates were matched across time. The Fano factor was then computed from the residual spikes.

*Network model:* We implemented a network model of our data in Python using the Brian simulator version 2. The network model was a biophysical network model of a randomly connected excitatory inhibitory (E-I) network with 1600 excitatory (E) and 400 inhibitory (I) leaky integrate-and-fire neurons. We used the Euler integration method with a time step of 0.1 ms. The network model was based on the model of an MT-like sensory circuit published by Wimmer and colleagues (Wimmer et al., 2015). All model equations and parameter values used were replicated from Wimmer (2015). Synaptic transmission mimicked AMPA and GABA_A_ receptor conductance dynamics. Excitatory neurons were divided into two populations, E1 and E2, each preferring opposite directions of motion. Connections within each population were stronger (w_*pref*_ = 1.3) compared to connections across populations (w_*null*_ = 0.7), mimicking the stronger coupling among cells with similar tuning. The two populations were then further divided into two equal sub-populations (E1i, E2i, E1o, E2o) that represented cells inside and outside the LPZ. We implemented two types of manipulations: the type of input into MT neurons (direction selective, non-direction selective, or no visual input) and changes in the balance of excitation and inhibition within MT.

The input into MT was modeled as a time-varying input current into each neuron:

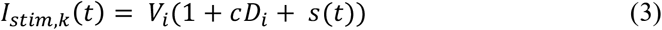

where *I_stim_* is the amount of current (nA) injected into each neuron (*k*) at over time (*t*). *V_i_* is the amount of visual input for a stimulus with no motion coherence (0.04 nA). *D_i_* is the amount of additional direction selective input to a neuron when the stimulus is moving in the preferred direction and is modulated by the motion coherence (*c*) of the stimulus. Time varying modulations in sensory input by the specific realization of the random dot stimulus are captured by *s(t)* where:

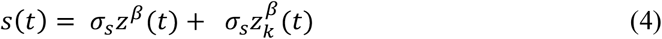

where *z^β^*(*t*) and 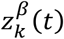 are independent Ornstein-Uhlenbeck processes with zero mean, s.d. equal to one, and a time constant of 20 ms. The term *z^β^*(*t*) represnts the ‘common’ part of the stimulus that is consistent across each neural population *β* and 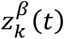 represents the “private” part of the stimulus that is unique to each neuron in each population. The amplitude of the temporal modulation of the stimulus is set by *σ_s_* (0.212). This allowed us to modulate the parameters of the stimulus into the populations of neurons “inside” and “outside” the LPZ to mimic different types of inputs into the LPZ in the absence of V1 input.

We also modified the E-I balance of the network by manipulating the probability of synaptic connections between neurons. Under balanced conditions, all neural connections (EE, EI, II) were connected with equal probability (p = 0.2). Under imbalanced conditions, the probability of connection between excitatory neuron outside the LPZ and inside the LPZ was increased (p = 0.3) and the probability of connection with inhibitory neurons inside the LPZ was decreased (p = 0.1). Finally, we reduced the weighting of synaptic inputs between excitatory neurons inside and outside the LPZ (w = 1.1). As in the experimental conditions, the “stimulus” lasted for 500 ms with a 1000 ms interval between repetitions. We tested network responses to coherences (0, 0.25, 5, 0.75, 1) in the preferred and null directions of each population.

### Histology

At the end of the recordings, the animals were given an intravenous lethal dose of sodium pentobarbitone and, following cardiac arrest, were perfused with 0.9% saline, followed by 4% paraformaldehyde in 0.1M phosphate buffer pH7.4. The brain was postfixed for approximately 24 hours in the same solution, and then cryo-protected with fixative solutions containing 10%, 20%, and 30% sucrose. The brains were then frozen and sectioned into 40 μm coronal slices. Alternate series were stained for Nissl substance and myelin (Gallyas, 1979). The location of recording sites was reconstructed by identifying electrode tracks, depth readings recorded during the experiment, and by electrolytic lesions performed at the end of penetrations. Area MT was defined by heavy myelination (Rosa and Elston, 1998). Recording sites were projected onto layer 4 lateral view reconstructions of MT (Fig. 2). All neurons reported here were histologically confirmed to be in MT including those in the two control animals.

*Estimation of LPZ:* To assess the changes in responses of in MT of V1-damaged animals compared with control animals, it was necessary to estimate the portion of MT that originally represented the same part of the visual field as the portion of V1 affected by the lesion. This portion of MT is referred to as the LPZ, as neurons inside the lesion projection zone no longer receive normal retinotopic inputs from V1 after the lesion. To establish the LPZ, isoeccentricity and polar lines were projected onto the lateral view reconstructions of MT in each individual case based on the average visuotopic map of this area (Fig 1D-G; (Rosa and Elston, 1998)). This average map was based on reconstructions of MT in six normal cases, which were superimposed after having been aligned and scaled to equal area. The visuotopy of MT has been demonstrated to be highly consistent across individuals (Rosa and Elston, 1998). The estimates of the pre-lesion visuotopic maps thus obtained were then fine-tuned for each case based on the size and extent of MT measured in each case (see above). Based on the visuotopic coordinates, the scotoma for each individual animal was then projected onto the lateral view reconstruction of MT (Fig 1D-G), thus defining a lesion projection zone for each individual case.

## Acknowledgements

This project was funded by the Australian Research Council (DE130100493 to LL; CE140100007 to MR) and by the National Health and Medical Research Council of Australia (GNT1066232 to LL and GNT1083152 to MR). We thank Katrina Worthy for the histological work. We also thank Janssen-Cilag for the donation of sufentanil citrate. KRH was partially supported by an Unrestricted Grant from the Research to Prevent Blindness Foundation to the Flaum Eye Institute at the University of Rochester.

## References

Ajina S, Pestilli F, Rokem A, Kennard C, Bridge H (2015) Human blindsight is mediated by an intact geniculo-extrastriate pathway Brown EN, ed. Elife 41–23. http://doi.org/10.7554/eLife.08935.001

Alexander I, Cowey A (2008) The cortical basis of global motion detection in blindsight. Exp Brain Res 192:407–411. http://doi.org/10.1007/s00221-008-15084

Atapour N, Worthy KH, Lui LL, Yu H-H, Rosa MGP (2017) Neuronal degeneration in the dorsal lateral geniculate nucleus following lesions of primary visual cortex: comparison of young adult and geriatric marmoset monkeys. Brain Struct Funct 222:3283–3293. http://doi.org/10.1007/s00429017-14044

Azzopardi P, Cowey A (2001) Motion discrimination in cortically blind patients. Brain 124:30–46.

Azzopardi P, Fallah M, Gross CG, Rodman HR (2003) Response latencies of neurons in visual areas MT and MST of monkeys with striate cortex lesions. Neuropsychologia 41:1738–1756. http://doi.org/10.1016/S0028–3932(03)00176–3

Barbur JL, Watson J, Frackowiak R, Zeki S (1993) Conscious visual perception without VI. Brain 116:1293–1302.

Barnes SJ, Franzoni E, Jacobsen RI, Erdelyi F, Szabo G, Clopath C, Keller GB, Keck T (2017) Deprivation-induced homeostatic spine scaling in vivo is localized to dendritic branches that have undergone recent spine loss. Neuron 96:871–882. http://doi.org/10.1016/j.neuron.2017.09.052

Berman RA, Wurtz RH (2010) Functional identification of a pulvinar path from superior colliculus to cortical area MT. J Neurosci 30:6342–6354. http://doi.org/10.1523/JNEUROSCI.6176-09.2010

Berman RA, Wurtz RH (2011) Signals conveyed in the pulvinar pathway from superior colliculus to cortical area MT. J Neurosci 31:373–384. http://doi.org/10.1523/JNEUROSCI.4738-10.2011

Brainard DH (1997) The Psychophysics Toolbox. Spat Vis 10:433–436.

Bridge H, Hicks SL, Xie J, Okell TW, Mannan S, Alexander I, Cowey A, Kennard C (2010) Visual activation of extra-striate cortex in the absence of V1 activation. Neuropsychologia 48:4148–4154. http://doi.org/10.1016/j.neuropsychologia.2010.10.022

Bridge H, Thomas O, Jbabdi S, Cowey A (2008) Changes in connectivity after visual cortical brain damage underlie altered visual function. Brain 131:1433–1444. http://doi.org/10.1093/brain/awn063

Britten KH, Shadlen MN, Newsome WT, Movshon JA (1992) The analysis of visual motion: a comparison of neuronal and psychophysical performance. J Neurosci 12:4745–4765.

Cavanaugh MR, Huxlin KR (2017) Visual discrimination training improves Humphrey perimetry in chronic cortically induced blindness. Neurology 88:1856–1864. http://doi.org/10.1212/WNL.0000000000003921

Cavanaugh MR, Zhang R, Melnick MD, Das A, Roberts M, Tadin D, Carrasco M, Huxlin KR (2015) Visual recovery in cortical blindness is limited by high internal noise. J Vis 15:9. http://doi.org/10.1167/15.10.9

Celeghin A, Diano M, de Gelder B, Weiskrantz L, Marzi CA, Tamietto M (2017) Intact hemisphere and corpus callosum compensate for visuomotor functions after early visual cortex damage. Proc Natl Acad Sci USA. http://doi.org/10.1073/pnas.1714801114

Chaplin TA, Allitt BJ, Hagan MA, Price NSC, Rajan R, Rosa MGP, Lui LL (2017) Sensitivity of neurons in the middle temporal area of marmoset monkeys to random dot motion. J Neurophysiol 118:1567– 1580. http://doi.org/10.1152/jn.00065.2017

Chaplin TA, Yu HH, Soares JGM, Gattass R, Rosa MGP (2013) A conserved pattern of differential expansion of cortical areas in simian primates. J Neurosci 33:15120–15125. http://doi.org/10.1523/JNEUROSCI.2909-13.2013

Cheong SK, Tailby C, Solomon SG, Martin PR (2013) Cortical-like receptive fields in the lateral geniculate nucleus of marmoset monkeys. J Neurosci 33:6864–6876. http://doi.org/10.1523/JNEUROSCI.520812.2013

Churchland MM et al. (2010) Stimulus onset quenches neural variability: a widespread cortical phenomenon. Nat Neurosci 13:369–378. http://doi.org/10.1038/nn.2501

Collins CE, Lyon DC, Kaas JH (2003) Responses of neurons in the middle temporal visual area after long-standing lesions of the primary visual cortex in adult new world monkeys. J Neurosci 23:2251–2264.

Darian-Smith C, Gilbert CD (1994) Axonal sprouting accompanies functional reorganization in adult cat striate cortex. Nature 368:737–740.

Das A, Tadin D, Huxlin KR (2014) Beyond blindsight: properties of visual relearning in cortically blind fields. J Neurosci 34:11652–11664. http://doi.org/10.1523/JNEUROSCI.1076-14.2014

Eysel UT, Schweigart G (1999) Increased receptive field size in the surround of chronic lesions in the adult cat visual cortex. Cereb Cortex 9:101–109.

Eysel UT, Schweigart G, Mittmann T, Eyding D, Qu Y, Vandesande F, Orban G, Arckens L (1999) Reorganization in the visual cortex after retinal and cortical damage. Restor Neurol Neurosci 15:153– 164.

Fritsches KA, Rosa MGP (1996) Visuotopic organisation of striate cortex in the marmoset monkey (Callithrix jacchus). J Comp Neurol 372:264–282.

Gallyas F (1979) Silver staining of myelin by means of physical development. Neurol Res 1:203–209.

Giannikopoulos DV, Eysel UT (2006) Dynamics and specificity of cortical map reorganization after retinal lesions. Proc Natl Acad Sci USA 103:10805–10810. http://doi.org/10.1073/pnas.0604539103

Girard P, Salin PA, Bullier J (1992) Response selectivity of neurons in area MT of the macaque monkey during reversible inactivation of area V1. J Neurophysiol 67:1437–1446.

Horton JC, Hoyt WF (1991) The representation of the visual field in human striate cortex. A revision of the classic Holmes map. Arch Ophthalmol 109:816–824.

Huxlin KR, Martin T, Kelly K, Riley M, Friedman DI, Burgin WS, Hayhoe M (2009) Perceptual relearning of complex visual motion after V1 damage in humans. J Neurosci 29:3981–3991. http://doi.org/10.1523/JNEUROSCI.4882-08.2009

Keck T, Scheuss V, Jacobsen RI, Wierenga CJ, Eysel UT, Bonhoeffer T, Hübener M (2011) Loss of sensory input causes rapid structural changes of inhibitory neurons in adult mouse visual cortex. Neuron 71:869–882. http://doi.org/10.1016/j.neuron.2011.06.034

Klüver H (1936) An analysis of the effects of the removal of the occipital lobes in monkeys. J Psychol: Interdiscip Appl 2:49–61. http://doi.org/10.1080/00223980.1941.9917017

Klüver H (1941) Visual functions after removal of the occipital lobes. J Psychol: Interdiscip Appl 11:23–45.

Lister WT, Holmes G (1916) Disturbances of vision from cerebral lesions, with special reference to the cortical representation of the macula. Proc R Soc Med 9:57–96.

Litwin-Kumar A, Doiron B (2012) Slow dynamics and high variability in balanced cortical networks with clustered connections. Nat Neurosci 15:1498–1505. http://doi.org/10.1038/nn.3220

Movshon JA, Newsome WT (1996) Visual response properties of striate cortical neurons projecting to area MT in macaque monkeys. J Neurosci 16:7733–7741.

Palagina G, Eysel UT, Jancke D (2009) Strengthening of lateral activation in adult rat visual cortex after retinal lesions captured with voltage-sensitive dye imaging in vivo. Proc Natl Acad Sci USA 106:8743–8747. http://doi.org/10.1073/pnas.0900068106

Pelli DG (1997) The VideoToolbox software for visual psychophysics: transforming numbers into movies. Spat Vis 10:437–442.

Poppel E, Held R, Frost D (1973) Residual visual function after brain wounds involving the central visual pathways in man. Nature 243:295–296.

Raninen A, Vanni S, Hyvarinen L, Nasanen R (2007) Temporal sensitivity in a hemianopic visual field can be improved by long-term training using flicker stimulation. J Neurol Neurosurg Psychiatry 78:66–73. http://doi.org/10.1136/jnnp.2006.099366

Riddoch G (1917) Dissociation of visual perceptions due to occipital injuries, with especial reference to appreciation of movement. Brain 40:15–57.

Ringach DL, Shapley RM, Hawken MJ (2002) Orientation selectivity in macaque V1: diversity and laminar dependence. J Neurosci 22:5639–5651.

Rodman HR, Gross CG, Albright TD (1989) Afferent basis of visual response properties in area MT of the macaque. I. Effects of striate cortex removal. J Neurosci 9:2033–2050.

Rodman HR, Gross CG, Albright TD (1990) Afferent basis of visual response properties in area MT of the macaque. II. Effects of superior colliculus removal. J Neurosci 10:1154–1164.

Rosa MGP, Elston GN (1998) Visuotopic organisation and neuronal response selectivity for direction of motion in visual areas of the caudal temporal lobe of the marmoset monkey (Callithrix jacchus): middle temporal area, middle temporal crescent, and surrounding cortex. J Comp Neurol 393:505–527.

Rosa MGP, Tweedale R, Elston GN (2000) Visual responses of neurons in the middle temporal area of new world monkeys after lesions of striate cortex. J Neurosci 20:5552–5563.

Sahraie A, Hibbard PB, Trevethan CT, Ritchie KL, Weiskrantz L (2010) Consciousness of the first order in blindsight. Proc Natl Acad Sci USA 107:21217–21222. http://doi.org/10.1073/pnas.1015652107

Sahraie A, Trevethan CT, MacLeod MJ, Murray AD, Olson JA, Weiskrantz L (2006) Increased sensitivity after repeated stimulation of residual spatial channels in blindsight. Proc Natl Acad Sci USA 103:14971–14976. http://doi.org/10.1073/pnas.0607073103

Sanders MD, Warrington EK, Marshall J, Wieskrantz L (1974) “Blindsight”: Vision in a field defect. Lancet 1:707–708.

Schmid LM, Rosa MGP, Calford MB, Ambler JS (1996) Visuotopic reorganization in the primary visual cortex of adult cats following monocular and binocular retinal lesions. Cereb Cortex 6:388–405.

Schmid MC, Mrowka SW, Turchi J, Saunders RC, Wilke M, Peters AJ, Ye FQ, Leopold DA (2011) Blindsight depends on the lateral geniculate nucleus. Nature 466:373–377.

Sincich LC, Park KF, Wohlgemuth MJ, Horton JC (2004) Bypassing V1: a direct geniculate input to area MT. Nat Neurosci 7:1123–1128. http://doi.org/10.1038/nn1318

Solomon SG, Tailby C, Cheong SK, Camp AJ (2010) Linear and nonlinear contributions to the visual sensitivity of neurons in primate lateral geniculate nucleus. J Neurophysiol 104:1884–1898. http://doi.org/10.1152/jn.01118.2009

Warner CE, Goldshmit Y, Bourne JA (2010) Retinal afferents synapse with relay cells targeting the middle temporal area in the pulvinar and lateral geniculate nuclei. Front Neuroanat 4:8. http://doi.org/10.3389/neuro.05.008.2010

Warner CE, Kwan WC, Bourne JA (2012) The early maturation of visual cortical area MT is dependent on input from the retinorecipient medial portion of the inferior pulvinar. J Neurosci 32:17073–17085. http://doi.org/10.1523/JNEUROSCI.3269-12.2012

Warner CE, Kwan WC, Wright D, Johnston LA, Egan GF, Bourne JA (2015) Preservation of vision by the pulvinar following early-life primary visual cortex lesions. Curr Biol 25:424–434. http://doi.org/10.1016/j.cub.2014.12.028

Weiskrantz L (1996) Blindsight revisited. Curr Opin Neurobiol 6:215–220.

Weiskrantz L, Warrington EK, Sanders MD, Marshall J (1974) Visual capacity in the hemianopic field following a restricted occipital ablation. Brain 97:709–728. http://doi.org/10.1093/brain/97.1.709

White AJ, Solomon SG, Martin PR (2001) Spatial properties of koniocellular cells in the lateral geniculate nucleus of the marmoset Callithrix jacchus. J Physiol 533:519–535.

Wimmer K, Compte A, Roxin A, Peixoto D, Renart A, la Rocha de J (2015) Sensory integration dynamics in a hierarchical network explains choice probabilities in cortical area MT. Nat Commun 6:6177. http://doi.org/10.1038/ncomms7177

Yamahachi H, Marik SA, McManus JNJ, Denk W, Gilbert CD (2009) Rapid axonal sprouting and pruning accompany functional reorganization in primary visual cortex. Neuron 64:719–729. http://doi.org/10.1016/j.neuron.2009.11.026

Yu H-H, Rosa MGP (2010) A simple method for creating wide-field visual stimulus for electrophysiology: mapping and analyzing receptive fields using a hemispheric display. J Vis 10:15–15. http://doi.org/10.1167/10.14.15

Yu H-H, Verma R, Yang Y, Tibballs HA, Lui LL, Reser DH, Rosa MGP (2010) Spatial and temporal frequency tuning in striate cortex: functional uniformity and specializations related to receptive field eccentricity. Eur J Neurosci 31:1043–1062. http://doi.org/10.1167/10.14.15

Yu HH, Chaplin TA, Egan GW, Reser DH, Worthy KH, Rosa MGP (2013) Visually Evoked Responses in Extrastriate Area MT After Lesions of Striate Cortex in Early Life. J Neurosci 33:12479–12489. http://doi.org/10.1111/j.1460-9568.2010.07118.x

Zhang X, Kedar S, Lynn MJ, Newman NJ, Biousse V (2006) Natural history of homonymous hemianopia. Neurology 66:901–905. http://doi.org/10.1212/01.wnl.0000203338.54323.22

